# Ventral tegmental area astrocytes regulate drug-cue associations and drug intake

**DOI:** 10.64898/2026.07.20.739241

**Authors:** Brandon I. Garcia-Castañeda, Angelica R. Ramos, Ashley N. Miller, Leonor G. Cedillo, Zoe Kirchner, Idaira Oliva, Alexey A. Soshnev, Michael D. Scofield, James D Lechleiter, Matthew J. Wanat

**Affiliations:** Department of Neuroscience, Developmental and Regenerative Biology, University of Texas at San Antonio, San Antonio, TX, USA; Department of of Anesthesia and Perioperative Medicine, Medical University of South Carolina, Charleston, South Carolina; Department of Cell Systems and Anatomy, University of Texas Health Science Center at San Antonio, San Antonio, TX, USA

## Abstract

Cocaine use disorder remains a critical public health concern with limited treatment options. Astrocytes are increasingly recognized as active regulators of neurotransmission and are emerging as important contributors to addiction neurobiology. Here, we examined how chemogenetic activation of astrocytic Gq signaling within the ventral tegmental area (VTA) influences cocaine-associated behaviors in drug-naïve and cocaine-experienced rats. Using a subthreshold dose of cocaine in a conditioned place preference (CPP) paradigm, we found that VTA astrocyte Gq activation facilitated the acquisition of cocaine CPP in drug-naïve rats but suppressed the development of cocaine CPP in cocaine-experienced rats. Additionally, chemogenetic activation of VTA astrocyte Gq signaling suppressed voluntary cocaine intake in a self-administration paradigm. Together, these findings demonstrate that VTA astrocyte Gq signaling shapes cocaine-associated behaviors in a drug history-dependent manner, highlighting VTA astrocytes as a potential therapeutic target in cocaine use disorder.

## Introduction

Reward-seeking behaviors are fundamental to survival, directing organisms toward resources necessary for life. In substance use disorders, however, these normally adaptive processes become maladaptive, as drugs of abuse override neural systems that evolved to reinforce the pursuit of natural rewards [1]. In cocaine use disorder, this pathological engagement of reward circuitry is marked by heightened craving and the attribution of excessive motivational salience to drug-paired cues, which can promote drug seeking and intake [2–4]. These maladaptive processes can be modeled in rodents using preclinical assays such as conditioned place preference (CPP) tests and intra-venous drug self-administration paradigms. These models serve as well-validated approaches for examining the neurobiological mechanisms underlying cocaine use disorder [5–8].

The mesolimbic dopamine system is central to the expression of drug-dependent behaviors and establishment of cue-outcome associations [9]. Specifically, ventral tegmental area (VTA) dopamine neurons and their projections to the nucleus accumbens orchestrate reward-guided behaviors that are exploited in cocaine use disorder [10–13]. Drug induced adaptations in this circuit are thought to underlie various maladaptive drug dependent behaviors [14,15]. Notably, long-lasting behavioral adaptations following repeated drug exposure are accompanied by alterations in synaptic plasticity onto VTA dopamine neurons. These changes, including increased excitatory synaptic strength, enhance the responsiveness of VTA dopamine neurons to cocaine and cocaine-paired cues [16–29]. Consistent with this framework, disrupting VTA dopamine signaling is sufficient to attenuate drug-dependent behaviors such as cocaine

CPP and cocaine self-administration [30–34]. These findings establish the central role of VTA dopamine signaling in the expression and reinforcement of cocaine-dependent behaviors. Despite this foundational understanding, identifying upstream regulators of drug-dependent behaviors remains an important goal for the development of therapeutic targets.

While prior research has extensively focused on how drugs alter neuronal afferent inputs that modulate VTA dopamine neuron activity [35], the contribution of astrocytes within this circuit has recently come into focus. Once viewed as passive support cells, astrocytes are now recognized as integral components of the tripartite synapse, where they actively regulate neuronal function and undergo drug-dependent adaptations [36–43]. In particular, astrocytes respond to neurotransmitters that activate Gq signaling pathways, leading to changes in gliotransmission and transporter function [44–47]. Prior work on astrocytic regulation of drug-related behaviors has largely focused on the nucleus accumbens [48–55]. However, emerging evidence suggests that VTA astrocytes may also shape cocaine-associated neuroadaptations and behavioral outcomes [56–61]. To investigate how VTA astrocytes regulate drug-motivated behaviors, we employed chemogenetic activation of astrocytic Gq signaling within the VTA and assessed its influence on cocaine-associated behaviors. Specifically, we examined how Gq activation in VTA astrocytes impacted the acquisition of cocaine CPP in cocaine-naïve and cocaine-exposed rats, as well as cocaine intake in a self-administration paradigm.

## Methods

### Subjects and surgery

All procedures were approved by the University of Texas at San Antonio Institutional Animal Care and Use Committee. Adult (P60-65; weighing 200-300 g) male and female Sprague Dawley rats (Inotiv/Envigo or Charles River) were pair-housed upon arrival, maintained on a 12 hr light/dark cycle, and allowed ad libitum access to water and chow prior to surgery. Surgeries were performed under isoflurane anesthesia. For intracranial viral injection surgeries, rats received bilateral injections of AAV5-GFAP-hM3D(Gq)-mCherry (‘Gq-DREADD’; RRID: Addgene_50478) targeting the VTA (relative to bregma: -5.6 mm posterior; ± 0.5 mm lateral; -8.0 mm ventral). The virus was microinjected using a 10 µl Hamilton syringe mounted on a syringe injector pump at a rate of 100 nl/min for a total of 1 µl per site. To ensure complete and successful viral injection, the microinjection syringe needle was left in place for 10 mins after viral injection. Following intracranial injection surgery, rats were allowed to recover for 3 weeks to allow for viral expression and were single-housed for the entirety of the remaining experimentation. For rats undergoing cocaine self-administration experiments, intravenous jugular catheterization surgeries were performed 2 weeks after the intracranial viral injection surgery. Rats were then allowed to recover for at least 1 week before beginning cocaine-self administration experiments.

### Cocaine conditioned placed preference (CPP)

Rats underwent a 4-day CPP paradigm. On day 1 (pre-test) and 4 (post-test), rats were allowed to roam freely for 30 mins between the separate chambers of the 2-chamber CPP apparatus (Med Associates). For each of the two conditioning days (days 2 and 3), the rats underwent 30-minute morning and afternoon sessions where they were confined to only one of the chambers. The assignment of the saline- and cocaine-paired chamber was counterbalanced across animals. Saline injections (1 ml/kg i.p.) were administered during the morning sessions and cocaine injections (10 mg/kg i.p in saline) were administered in the afternoon session. Rats additionally received a vehicle injection (1% DMSO in 1 ml/kg saline) 30 min prior to saline injections in the morning sessions and either a vehicle injection or CNO injection (3 mg/kg in 1% DMSO) 30 prior to the cocaine injections in the afternoon sessions (See **Figs. 1C, 2C**). Video tracking of behavioral outputs (time spent per side, time mobile, and distance traveled) was collected using ANY-maze software. Cocaine CPP was quantified as the change in the time spent in the cocaine-paired side (*post-test – pre-test*) and as the difference score during the post-test (*time spent on the cocaine-paired side – time spent on the saline-paired side*).

**Figure 1:**
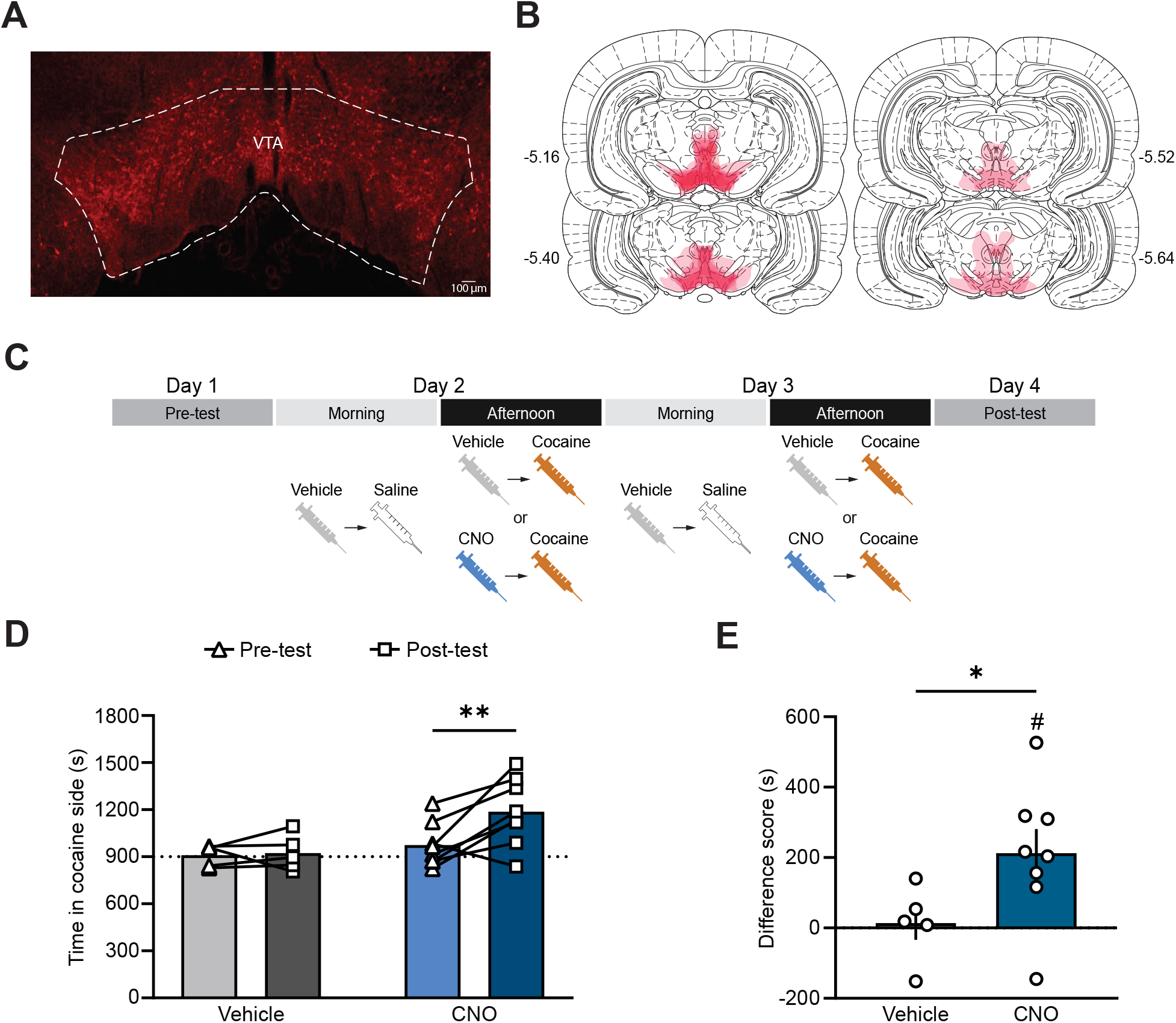
Chemogenetic activation of VTA astrocytes facilitates drug-cue learning in cocaine-naïve rats. (A) Representative Gq-DREADD viral expression in the VTA. (B) Gq-DREADD virus expression across subjects (vehicle: n = 5; CNO: n = 8). (C) Experimental timeline. (D) Cocaine CPP in vehicle- and CNO-treated rats (two-way ANOVA: effect of training F_(1,11)_ = 5.7, p = 0.036; post-hoc Šídák test: vehicle t_11_ = 0.19, p = 0.98; CNO t_11_ = 3.61, ** p = 0.0081). (E) Difference score of the time spent between the cocaine- and saline-paired chambers during the post-test session. The difference score was elevated in the CNO-treated group (one-sample t-test relative to zero: t_7_ = 3.1, # p = 0.017) and was significantly different when compared to the vehicle-treated group (Welch’s t test; t_10.93_ = 2.4, * p = 0.036).

### Cocaine self-administration training

Med Associates operant boxes were used for cocaine self-administration training sessions, including the previously described grid floors, house light, retractable levers, and cue lights above each lever [8,62]. To facilitate operant responding, rats were first trained to lever press for food rewards. Animals were placed on a mild food deprivation protocol to achieve 90% of their original body weight with an increase of 1.5% per week. Rats were trained to lever press for food pellet rewards (45 mg sucrose pellet) under a fixed ratio-1 (FR1) schedule with no timeout period after each successful press. After completing 100 successful lever presses within one 2-hr session, rats would next be trained to complete 60 trials under an FR1 schedule with a 20 s timeout period (2 sessions). Finally, rats would need to complete 2 more sessions but now they would be tethered to the i.v. catheter tubing. Rats would then be taken off food deprivation and initiate cocaine self-administration training.

Cocaine self-administration sessions under an FR1 reinforcement schedule (2 hours) were performed once daily for 18 total sessions with rats allowed ab libitum water and chow. Rats were first presented with the illumination of the house light and then the extension of both right and left levers. After completing a successful lever press to the active lever, the 20 s timeout period was initiated which coincided with the light above the active lever turning on, rats receiving a 0.3 mg/kg i.v. cocaine infusion, and the house light turning off. Rats were allowed to self-administer cocaine for 14 sessions. Rats that had a low level of responding (< 10 infusions) were excluded from the study. Rats received i.p injections of vehicle (1% DMSO in 1 ml/kg saline) or CNO (3 mg/kg in 1% DMSO) 30 min before sessions 15 or 17, with the order counterbalanced between subjects.

### Histology

Rats were intracardially perfused with 4% PFA, and brains were removed and postfixed for at least 24 h. Brains were subsequently placed in 15% and 30% sucrose solutions in PBS. Brains were then coronally sectioned using a Leica CM1860 Cryostat. For virus expression and location verification, immunohistochemistry was performed. Here, free-floating 40 µm coronal sections containing the VTA were blocked with a solution containing 0.3% Triton X-100 and 5% Normal Goat Serum (NGS, Invitrogen, Rockford, IL) in 0.1M PBS for 1 hour on an agitator at room temperature. Sections were then incubated at 4°C overnight with the primary antibody, mCherry (Abcam; ab13970; 1:500), diluted in 0.3% Triton X-100 and 5% NGS in 0.1M PBS. Slices were then washed 3 times for 10-minutes in 0.3% Triton X-100 in 0.1M PBS, then incubated at room temperature for 4 hours with the secondary antibody, Alexa Flour 594 (ThermoFisher; A-11042; 1:300) raised in goat, diluted in 0.3% Triton X-100 and 5% NGS in 0.1M PBS. The sections were then washed 3 times for 10 minutes in 0.1M PBS, mounted on SuperFrost Plus slides, and cover slipped with VectaShield Vibrance anti-fade mounting medium (Vector Laboratories, H-1700-10). Slides were then sealed with nail polish and stored in a dark area. The sections were then imaged using a Zeiss LSM 710 confocal microscope with a Zeiss Plan-Neofluar 5x objective and Zen imaging software. Images were collected by performing a tile scan of the sections at a frame size of 1024×1024 while keeping the pinhole size, laser power, and gain constant across images.

### Statistical analysis

Statistical analyses were performed using GraphPad Prism. A repeated measures ANOVA, paired t-tests, unpaired t-tests, and one sample t-tests were used in this study. In instances where animals did not complete the full training paradigm (due to experimenter error or other technical issues), a mixed-effects model was applied. Post-hoc significance was assessed with a Sidak’s multiple comparison test. Significance was set to α = 0.05 for all tests. All data are graphed as mean ± SEM. All statistical analyses are presented in **Supplementary Table 1**.

## Results

### Chemogenetic activation of VTA astrocytes facilitates cocaine-cue learning in drug-naïve rats

In our first set of experiments, we examined if stimulating Gq signaling in VTA astrocytes affects the acquisition of cocaine CPP. Cocaine CPP depends on elevating dopamine levels in the nucleus accumbens [63]. Prior research demonstrated that Gq-DREADD activation of astrocytes in the nucleus accumbens promotes the gliotransmission of glutamate [46]. Accordingly, we hypothesized that Gq-DREADD activation in VTA astrocytes would excite dopamine neurons and facilitate the acquisition of CPP to a subthreshold dose of cocaine.

Rats underwent surgery for bilateral virus injections targeting the VTA (**Fig. 1A-B**). Drug-naïve rats underwent a four-day CPP paradigm (**Fig. 1C**) using a subthreshold dose of cocaine (10 mg/kg) [64]. This subthreshold dose was selected to provide sufficient sensitivity to detect a potential facilitation in the development of cocaine CPP arising from Gq-DREADD activation in VTA astrocytes. Rats received two i.p. injections prior to conditioning sessions. In the morning sessions rats first received a vehicle injection (1% DMSO in 1 ml/kg saline) and were returned to their homecage. After 30 mins rats received an injection of saline and were placed in the conditioning chamber. In the afternoon sessions rats first received either a vehicle injection (1% DMSO in 1 ml/kg saline) or CNO injection (3 mg/kg in 1% DMSO) and were returned to their homecage. After 30 mins rats received an injection of cocaine (10 mg/kg) and were placed in the other conditioning chamber.

Vehicle-treated drug-naïve rats showed no significant change in time spent on the cocaine-paired side between pre- and post-test, consistent with the subthreshold cocaine dose failing to induce CPP. In agreement with our predictions, CNO-treated rats exhibited a significant increase in time spent in the cocaine-paired chamber at post-test (**Fig. 1D**). We additionally examined the difference score of the time spent between the cocaine- and saline-paired chambers during the post-test session. The difference score was elevated in the CNO-treated group and was significantly different when compared to the vehicle treated group (**Fig. 1E**). All together, these data indicate that chemogenetic activation of VTA astrocytes is sufficient to facilitate CPP acquisition in drug-naïve animals when paired with a subthreshold dose of cocaine.

### Prior cocaine exposure reverses the effect of VTA astrocyte activation on drug-cue learning

Prior studies demonstrated that previous drug exposure can prime animals for enhanced cocaine CPP acquisition[64–66]. Based on these findings and the results from the drug-naïve animals, we hypothesized that both vehicle- and CNO-treated cocaine-exposed rats would exhibit more robust acquisition of cocaine-CPP. Prior cocaine exposure is known to produce neuroadaptations that alter drug-cue learning [17–29], and emerging evidence suggests that drug history reshapes astrocyte signaling in ways that drive synaptic and circuit-level plasticity [49–57,67]. We therefore examined whether prior cocaine exposure modifies the behavioral effect of VTA astrocyte Gq-DREADD activation (**Fig. 2A-B**). Rats in the cocaine-exposed group received five daily injections of the subthreshold dose of cocaine (10 mg/kg) before undergoing the CPP protocol (**Fig. 2C**). Consistent with the known sensitizing effects of repeated cocaine exposure [64–66], vehicle-treated cocaine-exposed rats demonstrated robust CPP for the cocaine-paired chamber (**Fig. 2D**). Remarkably, and counter to the initial prediction, CNO-treated cocaine-exposed rats failed to establish CPP altogether (**Fig. 2D**). In the cocaine-exposed animals, the difference score was elevated in the vehicle-treated group and was significantly different when compared to the CNO-treated group (**Fig. 2E**). Gq-DREADD activation in VTA astrocytes did not affect the development of cocaine CPP in animals that received daily saline injections prior to undergoing CPP (**Supplemental Fig. 1**). Together, these findings indicate that VTA astrocyte Gq-DREADD activation facilitates drug-cue associations in drug-naïve animals but disrupts them following prior cocaine exposure, suggesting that drug exposure history fundamentally shifts the functional impact of VTA astrocyte signaling on establishment of drug-cue associations.

**Figure 2:**
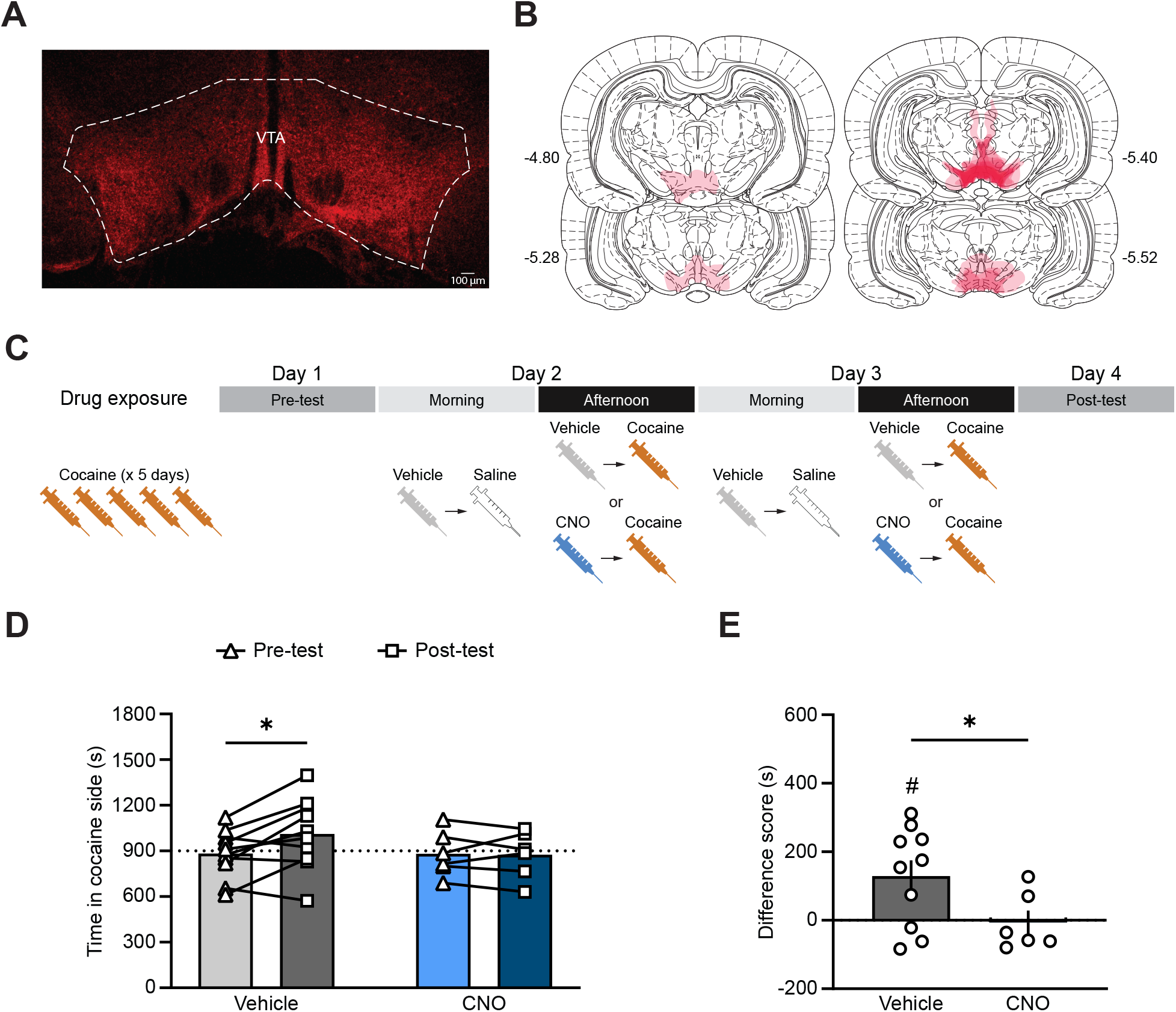
Chemogenetic activation of VTA astrocytes suppresses drug-cue learning in cocaine-exposed rats. (A) Representative Gq-DREADD viral expression in the VTA. (B) Gq-DREADD virus expression across subjects (vehicle: n = 10; CNO: n = 6). (C) Experimental timeline. (D) Cocaine CPP in vehicle- and CNO-treated rats that had prior cocaine exposure (two-way ANOVA: effect of drug F_(1,14)_ = 0.59, p = 0.045; post-hoc Šídák test: vehicle t_14_ = 3.24, * p = 0.011; CNO t_14_ = 0.12, p = 0.99). (E) Difference score of the time spent between the cocaine- and saline-paired chambers during the post-test session. The difference score was elevated in the vehicle-treated group (one-sample t-test relative to zero: t_9_ = 2.83, # p = 0.02) and was significantly different when compared to the CNO-treated group (Welch’s t test; t_14_ = 2.37, * p = 0.033).

### VTA astrocyte Gq-DREADD activation has differential effects on locomotor activity across drug history groups

Given the opposing effects of VTA astrocyte Gq-DREADD activation on cocaine-CPP acquisition in cocaine-naïve and cocaine-exposed rats (**Figs. 1-2**), we next examined whether these behavioral outcomes were accompanied by changes in locomotor activity during conditioning sessions. Cocaine exposure can produce behavioral sensitization, commonly measured as increased locomotor activity [27,66,68–70]. However, this effect is typically most robust at doses higher than the subthreshold dose used in these experiments [27,65]. Therefore, we analyzed time mobile and distance traveled during cocaine-paired conditioning sessions to determine whether locomotor activity paralleled the acquisition or disruption of cocaine CPP.

In drug-naïve rats, CNO-treated animals showed a significant decrease in distance traveled across cocaine-conditioning sessions. This effect appeared to be primarily driven by a reduction in locomotor activity during the day 3 conditioning session (**Fig. 3A**). There was also an overall reduction in the time mobile between conditioning days, with CNO-treated animals exhibiting lower levels of mobility relative to vehicle-treated animals in day 3 conditioning sessions (**Fig. 3B**), indicating a general decrease in mobility across conditioning rather than a CNO-specific locomotor enhancement. In contrast, cocaine-exposed rats showed no differences in either distance traveled (**Fig. 3C**) or time mobile (**Fig. 3D**) between vehicle- and CNO-treated groups. These findings indicate that VTA astrocyte Gq-DREADD activation does not mimic an additional cocaine-like stimulus or enhance cocaine-induced locomotor activity in a manner consistent with behavioral sensitization. Rather, the absence of an increase in locomotor activity suggests that opposing effects of VTA astrocyte activation on CPP acquisition are unlikely to be explained by nonspecific changes in cocaine-induced locomotor activity. Consistent with findings from previous literature [71–73], distance traveled did not correlate with CPP difference scores for the drug-naïve (r2 = −0.031, p = 0.60) or cocaine-exposed groups (r2 = −0.22, p = 0.17), suggesting that locomotor activity did not predict increased or decreased ability to make cocaine-cue associations.

**Figure 3:**
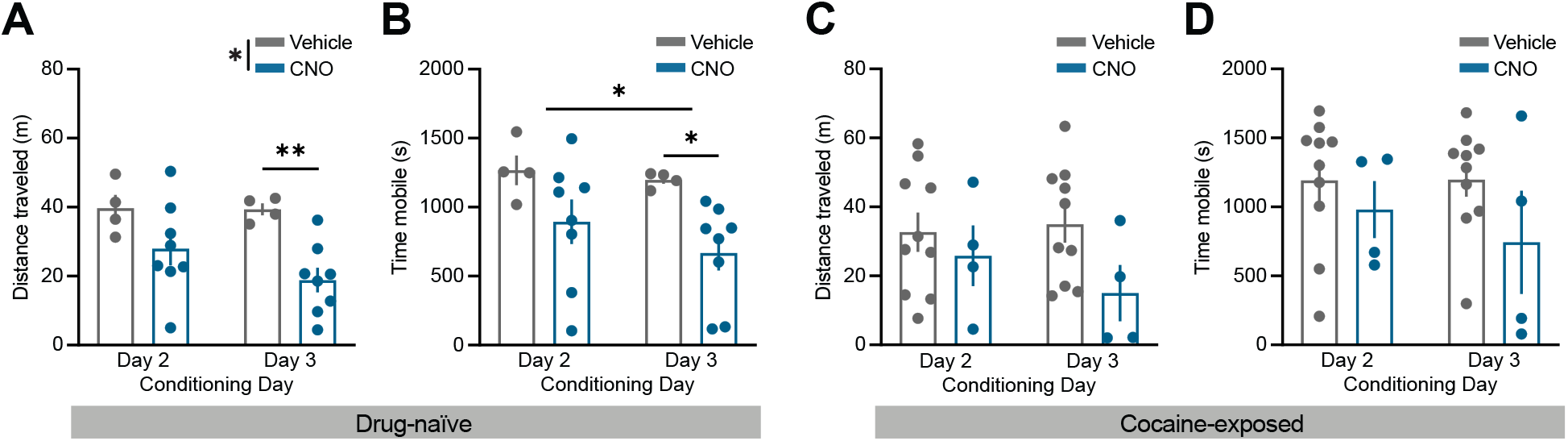
Effect of VTA astrocyte Gq-DREADD activation on locomotor activity. (A) There was a significant difference in the distance travelled during conditioning sessions between vehicle- and CNO-treated cocaine-naïve animals (mixed effects ANOVA: effect of treatment, F_(1,10)_ = 7.37, p = 0.022; post-hoc Šídák test for Day 3: t_20_ = 3.18, * p = 0.0095). (B) There was a significant decrease in the time mobile across conditioning sessions in drug-naïve rats (mixed-effects ANOVA: effect of training, F_(1,10)_ = 7.54, * p = 0.021; post-hoc Šídák test for Day 3: t_20_ = 2.43, * p = 0.049). (C) In rats with prior exposure to cocaine, there was no difference in the distance travelled during conditioning days relative between vehicle-injected rats and CNO-injected rats. (D) In rats with prior exposure to cocaine, there was no difference in the time mobile travelled during conditioning days relative between vehicle-injected rats and CNO-injected rats.

### Chemogenetic activation of VTA astrocyte Gq-signaling suppresses cocaine self-administration

Our CPP findings indicates that prior cocaine exposure induces adaptations in VTA astrocytes whereby stimulating Gq-signaling in VTA astrocytes can suppress the development of drug-cue associations. Given these findings, we next asked whether this effect extends to voluntary cocaine intake in an intravenous cocaine self-administration paradigm. Rats were implanted with jugular catheters and trained to self-administer cocaine across 14 consecutive 2-hour sessions (**Fig. 4A-B**), during which each active lever press delivered an intravenous infusion of cocaine (0.3mg/kg). On sessions 15 and 17, animals received counterbalanced i.p. injections CNO or vehicle 30 mins prior to the start of cocaine self-administration sessions. CNO injections significantly reduced the number of active lever presses for cocaine infusion relative to vehicle-injections sessions (**Fig. 4C**). These findings indicate that chemogenetic activation of Gq signaling in VTA astrocytes is sufficient to suppress voluntary cocaine intake following cocaine self-administration exposure.

**Figure 4:**
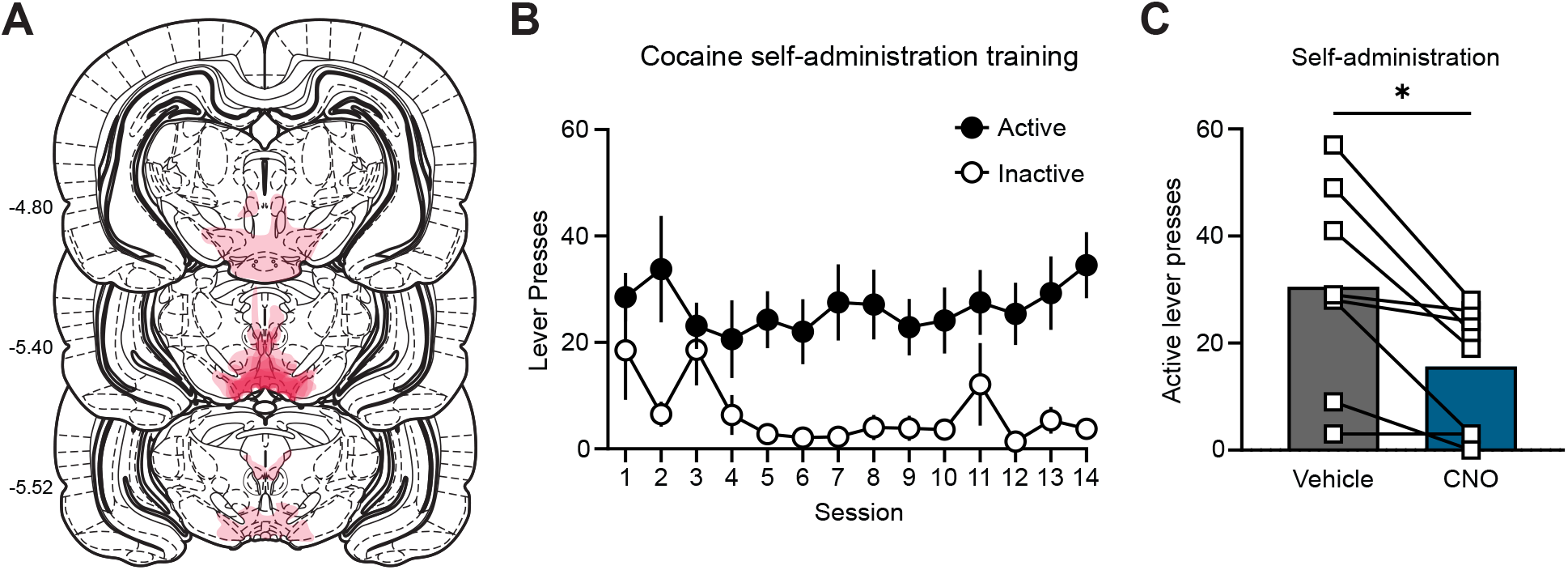
Chemogenetic activation of VTA astrocyte Gq-signaling suppresses cocaine self-administration. (A) Gq-DREADD virus expression across subjects (n = 8). (B) Average number of active and inactive lever presses during the first 14 cocaine self-administration session. (C) Rats received i.p. injections of vehicle or CNO prior to cocaine self-administration sessions 15 and 17, with the order counterbalanced across subjects. Rats performed significantly fewer active lever presses during the CNO injection sessions compared to vehicle injection sessions (paired t-test: t_7_ = 3.5, * p = 0.01).

## Discussion

Astrocytes are increasingly recognized as critical cellular elements that modulate how abused substances impact neuronal function and behavior [74]. Acute exposure to ethanol and psychostimulants increases astrocyte Ca2+ signaling both in culture and *in vivo* [51,75–79]. Ca2+ signaling in astrocytes regulate a host of cellular functions, including facilitating the release of transmitters [45]. Chemogenetic activation of Gq signaling in astrocytes enhances Ca2+ signaling and can promote the release of glutamate [46,80]. Accordingly, we hypothesized that Gq-DREADD activation in VTA astrocytes could functionally mimic the effect of acute drug exposure and facilitate the expression of dopamine-mediated behavior. To test this prediction, drug-naïve rats were trained on a cocaine CPP task using a subthreshold cocaine dose. We found that pairing Gq-DREADD activation of VTA astrocytes with a low dose cocaine injection resulted in rats developing cocaine CPP, whereas rats that only received the low dose cocaine injection during conditioning sessions failed to develop cocaine CPP. We additionally examined how Gq-DREADD activation affected motor activity during the cocaine conditioning sessions. Interestingly, pairing chemogenetic activation of VTA astrocytes with cocaine injections reduced motor activity relative to rats that only received the cocaine injections. While one might anticipate a higher level of activity in rats receiving cocaine injections with Gq-DREADD activation of VTA astrocytes, prior studies demonstrated that locomotor activity and drug-cue learning are not necessarily correlated [71–73]. Taken together, these data illustrate that stimulating Gq-signaling in VTA astrocytes facilitates the development of cocaine CPP in rats that have no prior history of exposure to the drug.

Our results highlight that Gq pathway activation in VTA astrocytes promotes the expression of a dopamine-dependent behavior (cocaine CPP) in rats without a prior history of drug exposure. One possibility is that this behavioral outcome arises from VTA astrocytes releasing excitatory molecules that preferentially act on VTA dopamine neurons. In support of this direct-excitatory model, Gq-DREADD activation in astrocytes has been shown to result in the release of a variety of excitatory molecules including glutamate, kynurenic acid, and ATP/ Adenosine [46,51,81]. Alternatively, astrocytes could indirectly disinhibit VTA dopamine neurons by dampening the activity of VTA GABA neurons. In support of the indirect-disinhibition model,

VTA astrocytes can inhibit VTA GABA neurons via the release of GABA through Swell1 volume-regulated anion channels [57]. Future mechanistic investigation of VTA astrocytes is necessary for identify how astrocytes regulate the development of cocaine CPP.

Prior studies demonstrated that previous drug exposure can facilitate the development of cocaine CPP [64–66]. We therefore predicted that prior cocaine exposure would (1) elicit cocaine CPP in rats trained with low dose cocaine injections and (2) produce even higher levels of cocaine CPP in rats trained with low dose cocaine injections paired with Gq-DREADD activation of VTA astrocytes. As expected, prior cocaine exposure produced cocaine CPP in rats trained using the subthreshold dosing procedure. However, we instead found no cocaine CPP in cocaine-exposed rats that received low dose cocaine injections coupled with Gq-pathway activation in VTA astrocytes. These data highlight how drug-dependent alterations in VTA astrocytes functionally switches the manner by which astrocytes regulate dopamine-dependent learning. Specifically, VTA astrocytes facilitate cocaine CPP in drug-naïve animals but suppress cocaine CPP in rats with prior cocaine experience.

Astrocytes undergo a variety of drug-mediated adaptations, though the majority of this research has focused on striatal astrocytes [39,42,48,55,74]. For example, following drug treatment striatal astrocytes can exhibit structural modifications in peripheral astrocyte processes [49,50], transcriptional changes [55], and alterations with the expression of glutamate transporters such as GLT-1 [58,82,83]. Future studies are needed to identify the specific cocaine-mediated alterations that occur within VTA astrocytes. However, one possibility is that drug treatment alters baseline activity and/or stimulus-elicited activity in VTA astrocytes. In support, optogenetic activation of VTA astrocytes using a 1Hz stimulation is aversive and is mediated by shunting the function of GLT-1 which selectively enhances glutamate transmission onto VTA GABA neurons that in turn suppressed dopamine neuron activity [58]. However, optogenetic activation of VTA astrocytes using a 10Hz stimulation is appetitive and is mediated by disinhibiting VTA GABA neurons via the glial-release of adenosine [61]. One must exercise caution about inferring biological relevance from using optogenetic manipulations in astrocytes that do not fire action potentials. However, these studies provide evidence that one can either elicit or suppress dopamine-dependent behavior depending upon the level of stimulation one applies to VTA astrocytes.

Given that Gq-DREADD activation in VTA astrocytes prevented the development of cocaine CPP in cocaine-exposed rats, we predicted this manipulation could additionally reduce voluntary drug intake in rats self-administering cocaine. Indeed, we found that acute activation of Gq signaling in VTA astrocytes suppressed cocaine intake in well-trained rats. Future experiments are needed to determine if chemogenetic activation of VTA astrocytes produces undesirable off-target effects on goal-directed behavior. However, we note that Gq-DREADD activation in VTA astrocytes did not affect motor activity in rats that had received prior cocaine treatment. Ultimately the net effect of abused substances on reward brain pathways depends on the acute and long-lasting changes on both neuronal and glial cellular populations. Our data though highlight that drug-mediated adaptations in VTA astrocytes may be exploited to reduce drug intake.

## Data Availability Statement

Data will be provided upon request.

## Author Contributions

BIG, ARR, IO, AAS, MDS, JDL, and MJW designed the experiments. BIG, ARR, ANM, LGC, and ZK, collected the data. BIG and ARR analyzed the data. BIG and MJW wrote the manuscript with input from the rest of the authors.

