## Supplemental Materials for "Ventral tegmental area astrocytes regulate drug-cue associations and drug intake"

### Supplemental Information:

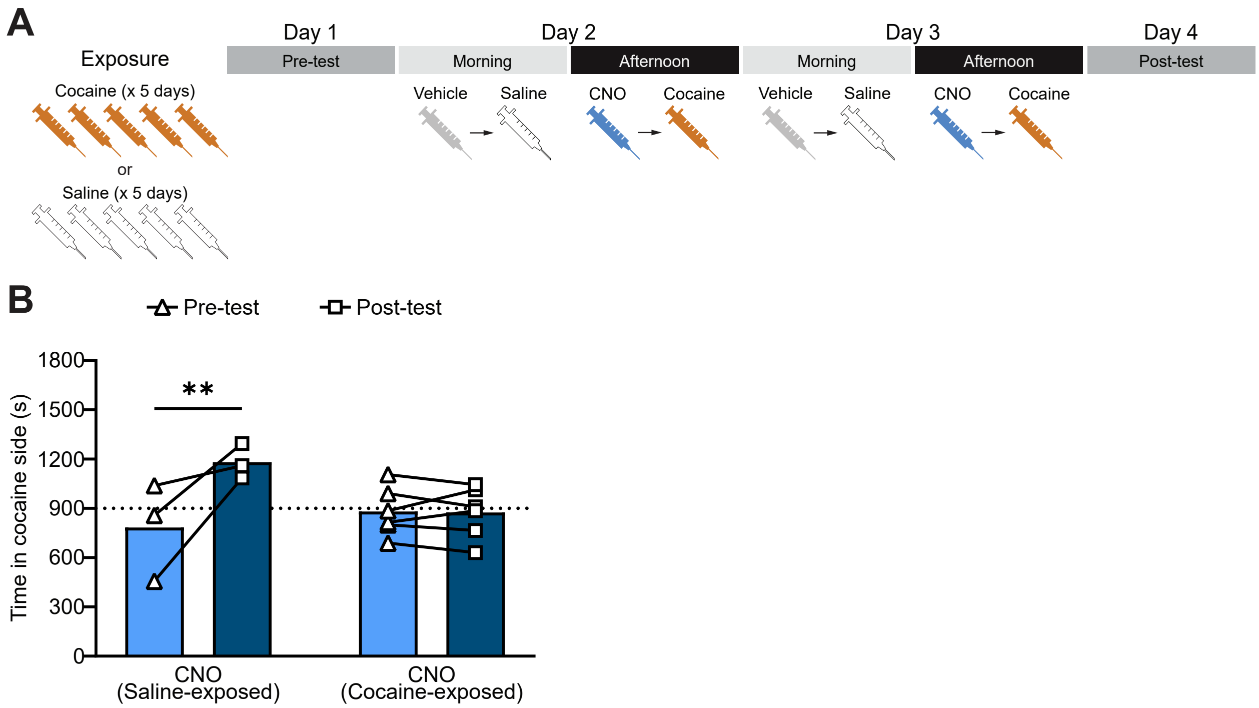


**Supplemental Figure 1: Chemogenetic activation of VTA astrocyte Gq-signaling elicits cocaine CPP in rats that had received prior injections of saline.** (A) Experimental timeline. (B) Cocaine CPP in CNO-treated rats that had prior daily injections of saline or cocaine (two-way ANOVA: effect of prior treatment $F_{(1,7)} = 12.82$, $p = 0.009$; effect of training $F_{(1,7)} = 13.66$, $p = 0.0077$; post-hoc Šídák test: saline-exposed $t_7 = 4.46$, ** $p = 0.0059$; cocaine-exposed $t_7 = 0.10$, $p = 0.99$).

### Supplementary Table 1

| Figure 1 | | | |
|---|---|---|---|
| all vehicle treated animals n=5 (4 female, 1 male), all CNO treated animals n=8 (5 female, 3 male) | | | |
| Panel D - Effect of VTA asrtocyte Gq-DREADD activation on cocaine CPP (pre-test vs post-test) | | | |
| Two-way ANOVA mixed effects model | Pre-Test vs Post Test | Vehicle vs CNO | Two-way interaction |
| | F (1, 11) = 5.6595, *p=0.0361 | F (1, 11) = 4.709, p=0.0528 | F (1, 11) = 4.396, p=0.0599 |
| Post-hoc Šídák's multiple comparisons test | Treatment | Adjusted P Value | t, df |
| | Vehicle | 0.9795 | t=0.1846, df=11 |
| | CNO | **0.0081 | t=3.614, df=11 |
| Panel E - Cocaine CPP difference score (vehicle vs CNO) | | | |
| Unpaired t test with Welch's correction | P Value (two tailed) | Welch-corrected t | Welch-corrected df |
| | *0.0355 | 2.397 | 10.93 |
| One sample t test | Treatment | P Value (two tailed) | Welch-corrected t, df |
| | Vehicle | 0.7863 | t=0.2899, df=4 |
| | CNO | *0.0168 | t=3.123, df=7 |

| Figure 2 | | | |
|---|---|---|---|
| all vehicle treated animals n=10 (4 female, 6 male), all CNO treated animals n=6 (2 female, 4 male) | | | |
| Panel D - Effect of VTA asrtocyte Gq-DREADD activation on cocaine CPP (pre-test vs post-test) | | | |
| Two-way ANOVA mixed effects model | Pre-Test vs Post Test | Vehicle vs CNO | Two-way interaction |
| | F (1, 14) = 3.551, p=0.0804 | F (1, 14) = 0.5924, *p=0.04543 | F (1, 14) = 4.319, p=0.0566 |
| Post-hoc Šídák's multiple comparisons test | Treatment | Adjusted P Value | t, df |
| | Vehicle | *0.0119 | t=3.236, df=14 |
| | CNO | 0.9908 | t=0.1226, df=14 |
| Panel E - Cocaine CPP difference score (vehicle vs CNO) | | | |
| Unpaired t test with Welch's correction | P Value (two tailed) | Welch-corrected t | Welch-corrected df |
| | *0.0328 | 2.368 | 14 |
| One sample t test | Treatment | P Value (two tailed) | Welch-corrected t, df |
| | Vehicle | *0.0197 | t=2.831, df=9 |
| | CNO | 0.8621 | t=0.1829, df=5 |

| Figure 3 | | | |
|---|---|---|---|
| all vehicle treated drug-naïve animals n=4 (4 female, 0 male), all CNO treated drug-naïve animals n=8 (5 female, 3 male), all vehicle treated drug-exposed animals n=10 (4 female, 6 male), all CNO treated drug-exposed animals n=4 (1 female, 3 male) | | | |
| Panel A - Drug-naïve distance traveled (m) cocaine-paired side | | | |
| Two-way ANOVA mixed effects model | Vehicle vs CNO | Day 2 vs Day 3 | Two-way interaction |
| | F (1, 10) = 7.369, *p=0.0218 | F (1, 10) = 3.483, p=0.0916 | F (1, 10) = 3.017, p=0.1130 |
| Post-hoc Šídák's multiple comparisons test | Day | Adjusted P Value | Adjusted P Value |
| | 2 | 0.16 | t=1.822, df=20 |
| | 3 | **0.0095 | t=3.177, df=20 |
| Panel B - Drug-naïve time mobile (s) on cocaine-paired side | | | |
| Two-way ANOVA mixed effects model | Vehicle vs CNO | Day 2 vs Day 3 | Two-way interaction |
| | F (1, 10) = 4.572, p=0.0582 | F (1, 10) = 7.543, *p=0.0206 | F (1, 10) = 2.086, p=0.1792 |
| Post-hoc Šídák's multiple comparisons test | Day | Adjusted P Value | t, df |
| | 2 | 0.1935 | t=1.714, df=20 |
| | 3 | *0.0487 | t=2.429, df=20 |
| Panel C - Drug-exposed distance traveled (m) cocaine-paired side | | | |
| Two-way ANOVA mixed effects model | Vehicle vs CNO | Day 2 vs Day 3 | Two-way interaction |
| | F (1, 12) = 2.139, p=0.1693 | F (1, 12) = 0.8698, p=0.3694 | F (1, 12) = 2.042, p=0.1785 |
| Post-hoc Šídák's multiple comparisons test | Day | Adjusted P Value | Adjusted P Value |
| | 2 | 0.7614 | t=0.6664, df=24 |
| | 3 | 0.1224 | t=1.948, df=24 |
| Panel D - Drug-exposed time mobile (s) on cocaine-paired side | | | |
| Two-way ANOVA mixed effects model | Vehicle vs CNO | Day 2 vs Day 3 | Two-way interaction |
| | F (1, 12) = 1.516, p=0.2419 | F (1, 12) = 1.532, p=0.2395 | F (1, 12) = 1.671, p=0.2205 |
| Post-hoc Šídák's multiple comparisons test | Day | Adjusted P Value | Adjusted P Value |
| | 2 | 0.7157 | t=0.7395, df=24 |
| | 3 | 0.2356 | t=1.587, df=24 |

| Figure 4 | | | |
|---|---|---|---|
| all animals n=8 (6 female, 2 male) | | | |
| Panel C - Effect of VTA asrtocyte Gq-DREADD activation on cocaine self-administration | | | |
| Paired t test | P Value (two tailed) | Paired t test t | Paired t test df |
| | *p=0.0101 | t=3.494 | df=7 |

| Supplementary Figure 1 | | | |
|---|---|---|---|
| all drug-exposed CNO treated animals n=6 (2 female, 4 male), all saline-exposed CNO treated animals n=3 (1 female, 2 male) | | | |
| Panel B - Effect of cocaine vs saline pre-treatment on VTA asrtocyte Gq-DREADD activation during cocaine CPP (pre-test vs post-test) | | | |
| Two-way ANOVA mixed effects model | Saline-exposed vs drug-exposed | pre-test vs post-test | Two-way interaction |
| | F (1, 7) = 12.95, *p=0.0088 | F (1, 7) = 13.63, *p=0.0077 | F (1, 8) = 0.5291, p=0.4877 |
| Post-hoc Šídák's multiple comparisons test | Treatment | Adjusted P Value | Adjusted P Value |
| | CNO saline-exposed | **0.0059 | t=4.456, df=7 |
| | CNO drug-exposed | 0.9941 | t=0.1003, df=7 |
